# Topographical changes in extracellular matrix during skin fibrosis and recovery can be evaluated using automated image analysis algorithms

**DOI:** 10.1101/2024.02.23.581810

**Authors:** Rachel H. Wyetzner, Ella X. Segal, Anna R. Jussila, Radhika P. Atit

**Author notes:** Correspondence: Mailing Address: Millis Science Center, 2074 Adelbert Road, Cleveland, Ohio, USA, 44106.

## Abstract

Skin fibrosis is characterized by fibroblast activation and intradermal fat loss, resulting in excess deposition and remodeling of dermal extracellular matrix (ECM). The topography of the dominant ECM proteins, such as collagens, can indicate skin stiffness and remains understudied in evaluating fibrotic skin. Here, we adapted two different unbiased image analysis algorithms to define collagen topography and alignment in a genetically inducible and reversible Wnt activation fibrosis model. We demonstrated that Wnt activated fibrotic skin has altered collagen fiber characteristics and a loss of collagen alignment, which were restored in the reversible model. This study highlights how unbiased algorithms can be used to analyze ECM topography, providing novel avenues to evaluate fibrotic skin onset, recovery, and treatment.

## Introduction

Fibrosis impacts one in four individuals globally and occurs in all soft tissues [1] (Fig. 1A). It is characterized by excessive deposition of extracellular matrix (ECM), typically resulting in a loss of organ architecture and function. There are currently no therapies to prevent or reverse existing fibrosis. However, there are many histomorphological similarities among fibrosis of each organ, such that new mechanistic insights in one organ system will have implications in others [2]. Skin is an optimal system for studying fibrosis because of its accessibility and distinct ECM rich dermis and dermal white adipose tissue (DWAT) layers. Skin fibrosis includes scarring, scleroderma, and keloids, among other clinical presentations [3]. During the onset of acute and chronic models of skin fibrosis, there is a thickening of the dermal layer and a depletion of the DWAT [4,5,6]. This dermal thickening involves fibroblast activation, resulting in increased ECM deposition [7,8].

**Fig. 1.**
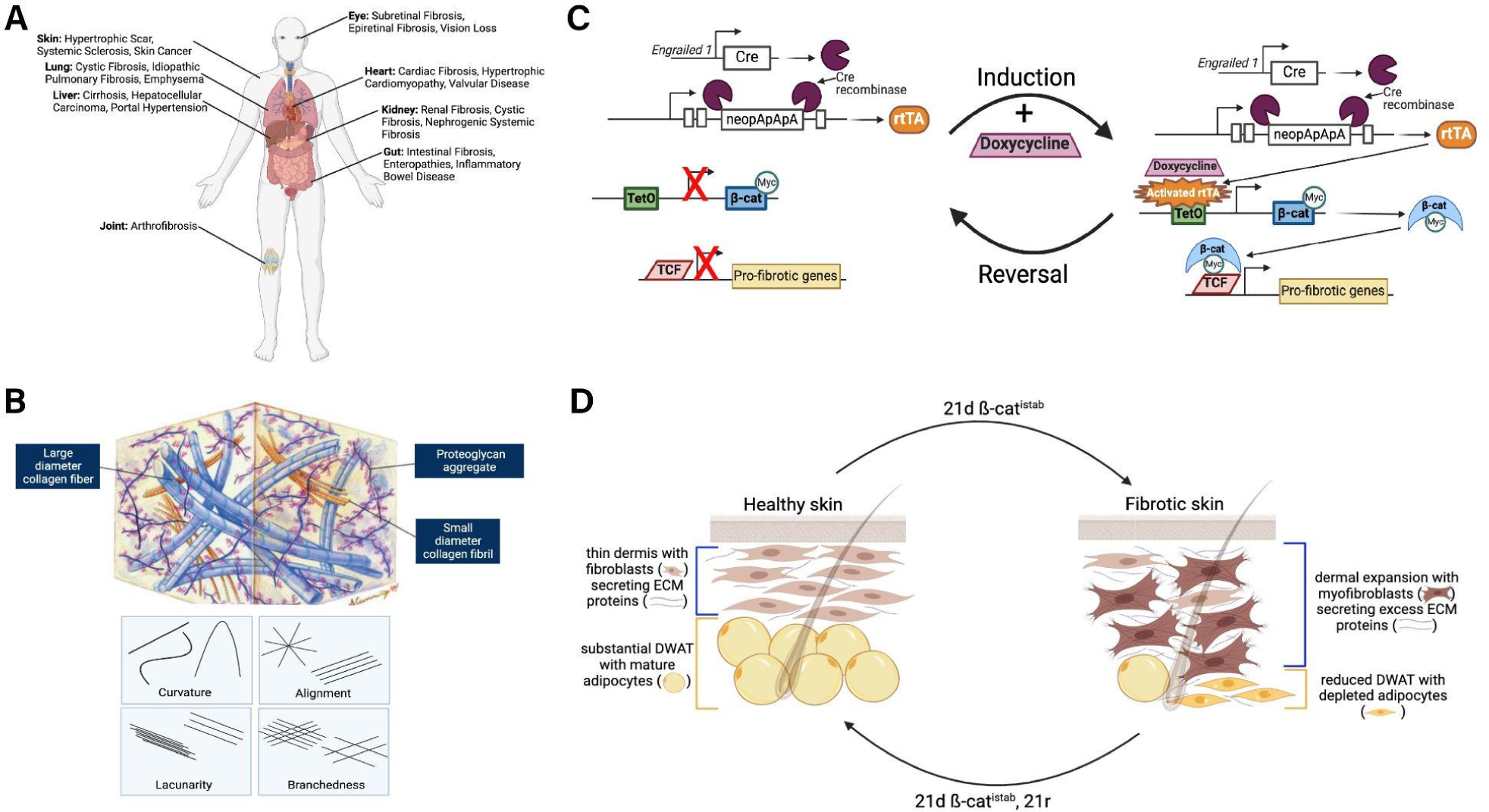
Types of surface topographies and applications in disease contexts. (A) Schematic illustrating the types and disease contexts of fibrosis in each organ. (B) Schematic depicting the extracellular matrix and different aspects of its topography. (C) Schematic depicting the genetic mouse lines in the inducible and reversible model of Wnt signaling induced fibrosis used in this study. (D) Schematic illustrating the difference in dermal and DWAT thickness observed during the induction and reversal of fibrosis using our Wnt activation model.

In the skin, the ECM is primarily composed of collagen fibrils, including collagens I and III. These fibrils are right-handed coils composed of three left-handed coiled polypeptide chains and arranged into bundles [9]. The orientation of collagen fibers is crucial for the mechanical properties of the skin. Collagen fibers typically have a largely compact and ordered organization, following a basketweave pattern to resist tensile stress in multiple directions [9,10]. Accumulated ECM in the dermis during chronic fibrosis is predominantly collagen. While the quantity of collagen is a commonly used metric when studying skin fibrosis, understanding the organization and quality of the deposited collagen in fibrotic skin is equally important for evaluating potential treatments. It is well-established that the mechanical stiffness of ECM is significantly elevated in fibrotic states, resulting in a positive feedback loop due to mechanoactivation of myofibroblasts [11]. Atomic force microscopy (AFM) indentation has been most used to characterize mechanical properties of collagen in the skin at the nanoscale, generating outputs such as the elastic modulus of the probed area [12,10]. However, AFM can only capture small areas of surface information from a histological section, resulting in an incomplete picture of the mechanical environment [13,14].

The best understood mechanism of regulating mechanical stiffness in the skin is by modifying the topography of collagen [15]. Topography refers to the surface features of a material or environment, such as curvature, branchedness, lacunarity, and alignment [16] (Fig. 1B). Collagen matrix cross linking, density, and fiber orientation are among the most crucial topographical characteristics involved in regulating mechanical stiffness [16]. Thus, in the absence of mechanical information from techniques such as AFM, we can use topographical information about ECM in healthy, fibrotic, and treated states to eventually make inferences about changes in ECM quality. Various automated softwares have been developed to measure topographical metrics of ECM, both at the tissue and individual fiber levels [17,18,19,20]. These softwares differ in the type of algorithm used, with the most common being fiber segmentation, image transformation, and curvelet-based algorithms [17]. Importantly, these softwares allow for acquisition of topographical information *in vivo* in a low-cost, efficient, and unbiased way.

In this study, we utilize a genetically inducible and reversible fibrotic murine model to study changes in collagen topography in the dermal ECM during the onset and reversal of fibrosis [4]. Given its simplicity and amenability to work on archived paraffin embedded tissue, we use picrosirius red (PSR) staining with polarized light microscopy for collagen detection in dorsal skin [21]. To analyze collagen at both the tissue and individual fiber level in an unbiased way, we employ two image analysis softwares: *Alignment by Fourier Transform (AFT)* and *The Workflow Of Matrix BioLogy Informatics (TWOMBLI)*, respectively. We utilize our recently developed model of inducible and reversible Wnt signaling activation in the skin to test if these algorithms can evaluate the topography of dermal collagen and demonstrate if skin fibrosis topography features are truly reversible. We found that there are several changes in dermal collagen topography induced by the onset of fibrosis that can be reversed upon removal of a fibrotic stimulus. Specifically, fibrotic changes in collagen topography include a decrease in collagen anisotropy and hyphal growth unit, yet increased matrix endpoints. Ultimately, this study demonstrates that unbiased algorithms can be used to analyze the topography of ECM in diseased tissues, providing insight for new ways to evaluate fibrosis treatment and therapeutic targets.

## Materials and Methods

### Mouse Handling and Lines

To study collagen topography collagen in dorsal skin under a fibrotic stimulus, *En1Cre* [22]; *Rosa26rtTA-EGFP* [22] (Jax Stock 005572); *TetO-deltaN89 β-catenin* [24] lines were used and genotyped. The significance of *En1Cre* lineage cells in skin has been established [28,29,30]. To induce *TetO-deltaN89 β-catenin-myc* tagged transgene expression in the *En1Cre*; *R26rtTA* recombined cells, 21-day old (p21) triple transgenic mice (β-cat^istab^) were given 2 mg/ml doxycycline in water (Sigma-Aldrich, St. Louis, MO) and 6 g/kg of dietary doxycycline in rodent chow (Envigo-Harlan) for three weeks [4]. To study collagen topography in dorsal skin during reversal from established fibrosis, Wnt induced fibrotic mice were subsequently switched from age p42 to p66 to a standard chow and water diet for three weeks (21d β-cat^istab^, 21r). At the analyzed time points, mice were euthanized and dorsal skin was processed for paraffin sections as detailed by Jussila *et al*., 2021 [4]. Litter-matched controls were studied for each experiment and at least two to four litters were used for phenotypic analysis. All animal experiments were approved by Case Western Reserve University Institutional Animal Care and Use Committee; all procedures were in accordance with AVMA guidelines (Protocol 2013-0156, approved 21 November 2019, Animal Welfare Assurance No. A3145-01).

### Histological Stains

To study the collagen topography in dorsal skin, dorsal mouse skin from p42 and p66 mice was isolated and drop-fixed in 10% neutral buffered formalin for 1 hour at 4°C [4]. The skin was then processed for paraffin sectioning and cut in the sagittal plane at 7 microns. Skin sections were placed parallel to each other on charged slides, with the epidermis side of each section perpendicular to the length of the slide (Supplementary Fig. 1A). PSR staining, which highlights the natural birefringence of collagen fibers when exposed to polarized light, was adapted from Gaytan *et al*., 2020 and performed on sections using commercial PSR stain (Electron Microscopy Sciences, 26357-02) [25].

### Image Analysis

For PSR-stained dorsal skin sections, uniformly sized Regions of Interest (ROIs) were taken from the lower dermis immediately above the DWAT in controls and from the remodeled DWAT above the panniculus carnosus muscle layer in mutants. Two ROIs were taken per image and 3 non-overlapping images per animal were included for analysis. Sections were positioned on the microscope stage so that the epidermis of each section was parallel to the top of the captured image. The images used for the main analysis were taken with the stage at 0 degrees. However, images were also taken after rotating the stage 45 degrees to determine differences in the fibers captured based on the angle of imaging, which have been observed by other groups [21].

#### Alignment by Fourier Transform (AFT)

To determine the alignment of collagen, all ROIs were converted to 16-bit grayscale using FIJI/ImageJ and input into *Alignment by Fourier Transform* (AFT) [17], using MATLAB (Mathworks, v2022a). The following parameters were used for calculating an image order parameter: Window Size: 30 pixels, Window Overlap: 50%, Neighborhood Radius: 2x vectors. Local masking and filtering were not applied to the images. Vector angle distribution within the ROI can be visualized as a heat map of vector orientations and quantified collectively as an order parameter. The order parameter can range between 0, for random alignment or isotropy, and 1, for perfect alignment or anisotropy. More homogeneous heat maps indicate a greater degree of anisotropy whereas more heterogeneous heat maps indicate a greater degree of isotropy. The median order parameters from twelve ROIs per animal were averaged and graphed.

#### The Workflow Of Matrix BioLogy Informatics (TWOMBLI)

To identify matrix patterns in the PSR-stained dorsal skin sections, the ROIs taken from these sections were run through TWOMBLI in Fiji/ImageJ [18] using the following parameters: Contrast Saturation: 0.35, Minimum Line Width: 5, Maximum Line Width: 10, Minimum Curvature Window: 20, Maximum Curvature Window: 70, Minimum Branch Length: 10, Maximum Display HDM: 220, Minimum Gap Diameter: 9. Line masks of the matrix network are generated via the ImageJ Ridge Detection tool. From these line masks, the following metrics were calculated per ROI: curvature, fractal dimension, number of branch points, total length, number of endpoints, lacunarity, and alignment. Averages of each metric were generated per animal from six total ROIs.

#### Principal Component Analysis (PCA)

A PCA biplot was generated for each skin fibrosis model control, mutant pair using the TWOMBLI output in RStudio. Each ROI was used as a datapoint so that there were more samples than variables being evaluated. Outliers were removed using the outlier calculator in Microsoft Excel and covariance between variables was checked using RStudio before running the PCA. Based on low covariance values, the curvature metrics were removed from the imported data. Principal components (PCs) were determined and PC1 and PC2 were used to generate a PCA biplot. The variance explained by each PC is displayed along the corresponding axis in the biplot.

### Statistical Analysis

Microsoft Excel version 16.16.27 for Mac (Microsoft) and GraphPad Prism version 9.0.0 for Mac (GraphPad Software) were used to produce all graphs and statistical analysis. Data are presented as mean ± SEM in all graphs. Outliers were excluded using the outlier calculator in Microsoft Excel. Because of the unequal variance and sample numbers, the pairwise sample comparisons for Collagen alignment and Collagen endpoints were performed using unpaired, two-tailed Welch’s t-test. p-values for statistical tests in all figures are represented using *=p<0.05, **=p<0.01, ***=p<0.001 and ****=p<0.0001.

## Results

### Fibrotic remodeling in skin results in a loss of collagen anisotropy in the newly remodeled DWAT

Understanding fibrotic changes in collagen topography can yield significant insight into fibrotic changes in collagen quality and cellular behavior, while also providing an avenue for evaluation of potential treatments. Using an inducible and reversible model of Wnt activation, we investigated the topography of collagen at the tissue and fiber level in the fibrotic and reversal state from established fibrosis. Canonical Wnt/β-catenin signaling is a conserved stimulus of tissue fibrosis in skin, as well as other organs [26,27]. The *Engrailed1 (En1)* gene is expressed in mouse embryonic fibro-adipogenic progenitors from E11.5 that differentiate into all derivatives of dorsal skin dermal fibroblasts and dermal adipocyte stem cells [28,29,7]. *Engrailed1*-lineage derived dermal fibroblasts are the key source of profibrotic fibroblasts in acute skin wounding models [30,7,31]. We therefore used a Wnt activation model that inducibly and reversibly expresses myc-tagged stabilized β-catenin (β-cat^istab^) in embryonic *Engrailed1* lineage-derived fibroblasts and adipocytes (Fig. 1C).

After three weeks of Wnt activation (21d β-cat^istab^) mice, we previously found a significant increase in dermal thickness and proportion of high density collagen matrix (HDM) as well as a decrease in DWAT thickness, as is characteristic of skin fibrosis (Fig. 1D, 2A, quantified in Jussila et al., 2021) [4]. Subsequently, after three weeks of reversal from Wnt activation in mutants (21d β-cat^istab^, 21r), we previously found that dermal thickness and proportion of HDM decreased while the DWAT tissue also recovered, suggesting a reversal from established fibrotic condition (Fig. 1D, 2B) [4]. However, changes in collagen topography during fibrosis onset and in this recent reversible model of skin fibrosis remains poorly understood [4].

**Fig. 2.**
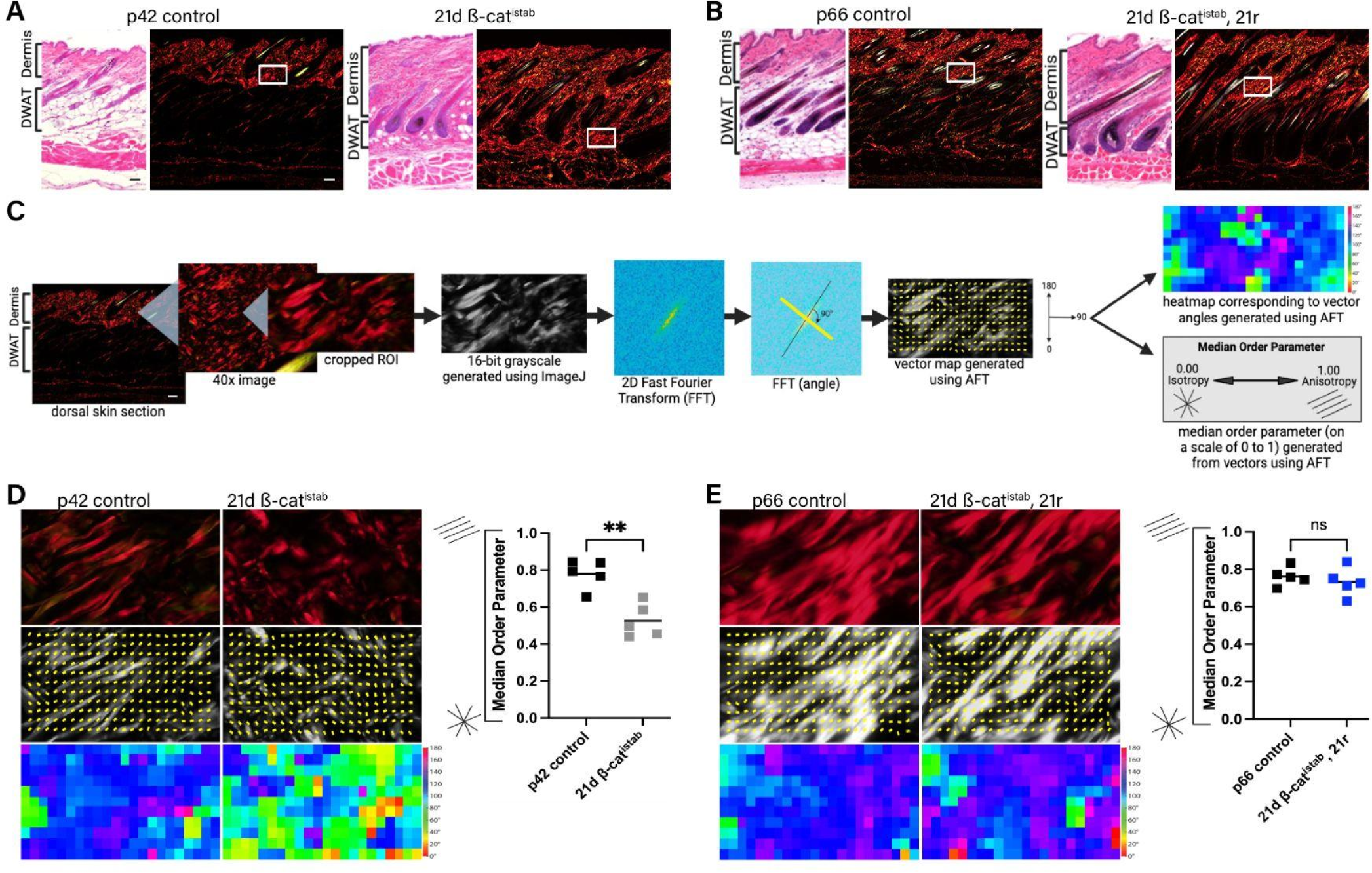
Isotropic organization of collagen is dependent on a sustained fibrotic stimulus. (A, B) H&E and picrosirius red-stained polarized light images of dorsal skin sections from fibrotic and rescued skin, as well as the corresponding controls. White boxes delineate regions considered for each ROI. (C) Schematic depicting the workflow for Alignment by Fourier Transform (AFT). (D,E) Cropped ROIs from 40x polarized light images of dorsal dermal collagen stained with picrosirius red (top row), and corresponding vector and heat maps showing the orientation of collagen (bottom two rows). n = 5 controls, 5 mutants in each figure. Median order parameters calculated from vector maps were plotted, with each data point representing the average from 6 ROIs obtained from 3 different sections per animal. Scale bar = 100 microns.

To determine the topography of collagen in fibrotic skin, sections of dorsal skin from 21d β-cat^istab^ mice and litter matched controls were stained with PSR followed by polarized light microscopy imaging [32]. Next, we analyzed the alignment of the fibers through adaptation of the *Alignment by Fourier Transform* (*AFT*) algorithm (Fig. 2C) [17]. Given that fibroblasts from the lower reticular dermis predominantly contribute to collagen synthesis during dermal repair, we focused our ROIs in the dermis adjacent to the DWAT layer in control skin [33]. In 21d β-cat^istab^ mutant skin, the DWAT layer becomes lipid depleted and undergoes extensive remodeling as previously shown by Collagen Hybridizing Peptide (CHP) staining [4]. We therefore placed ROIs in the newly remodeled DWAT. ROIs were run through *AFT*, which utilized a 2D Fast Fourier Transform to generate a vector map corresponding to the organization of collagen fibers (Fig. 2C) [17]. Then, a median order parameter was calculated from these vectors and a heat map corresponding to the orientation of the vectors was generated. The median order parameter ranged from 0 to 1, with 0 indicating complete isotropy, or disorganization, and 1 indicating complete anisotropy, or perfect alignment.

We found that collagen fibers were predominantly anisotropic in the healthy dermis. This was indicated by a high median order parameter and largely homogeneous heatmap outputted by the *AFT* software (Fig. 2D). However, following three weeks of Wnt activation, this collagen anisotropy was lost, as was demonstrated by a significantly lower median order parameter and more heterogeneous heatmap (Fig. 2D). In the 21d β-cat^istab^, 21r mutants, the anisotropy in collagen organization was recovered because the median order parameter and heatmap homogeneity was comparable to litter matched controls (Fig. 2E). These results indicated that fibrotic changes in collagen matrix alignment at the tissue level can be reversed and rescued upon eliminating a fibrotic stimulus or an established fibrotic state. Additionally, such changes in collagen matrix alignment that may not be distinguishable by eye in stained histological sections, (Fig. 2A) can be quantified by automated softwares (Fig. 2D, 2E).

### Fibrotic remodeling in skin results in changes in collagen matrix alignment, number of endpoints, and hyphal growth unit

In addition to alignment of collagen, we also aimed to determine how other aspects of collagen topography change during skin fibrosis. To test topographical characteristics of collagen fibers, we adapted *The Workflow Of Matrix BioLogy Informatics (TWOMBLI)* ImageJ macro plugin generated by the Sahai Lab [18]. TWOMBLI utilized the ridge detection tool in ImageJ to generate a line mask representing the ECM in an image [18]. From this mask, TWOMBLI calculated metrics of matrix alignment, length, branching, end points, gaps, fractal dimension, curvature, and the distribution of fiber thickness (Fig. 3A). We analyzed the previously described ROIs (above) with *TWOMBLI*. We then generated principal component analysis (PCA) biplots with the output matrix metrics to first determine if each set of mutants separated from its litter matched controls across these metrics and second, identify which metrics contributed most to any separation.

**Fig 3.**
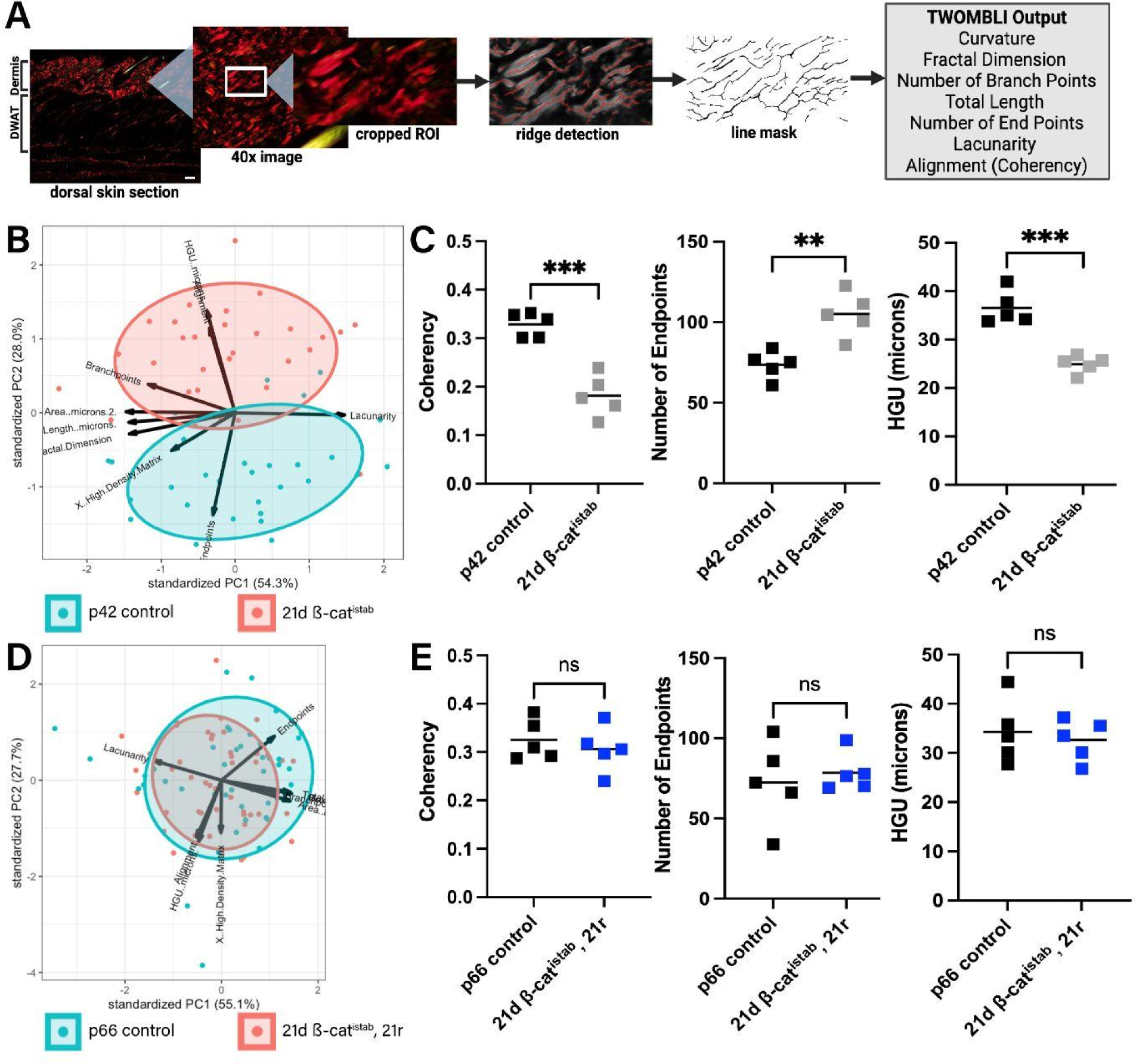
Collagen alignment, number of endpoints, and hyphal growth unit are dysregulated during fibrosis. (A) Schematic depicting workflow for TWOMBLI algorithm implemented in FIJI to quantify a broad range of matrix metrics. (B, D) Principal component analysis (PCA) plots showing the variance between each control/mutant pair and the TWOMBLI metrics that distinguish each control group and corresponding mutant group. Each data point represents one ROI and 6 ROIs were used for each animal. n=5 animals for each condition. (C, E) TWOMBLI output for coherency (0 = isotropy, 1 = anisotropy), number of endpoints, and hyphal growth unit for control/mutant pair. HGU: hyphal growth unit. Scale bar = 100 microns

We found that 21d β-cat^istab^ mice and litter matched controls separated greatly across *TWOMBLI* matrix metrics, with collagen alignment and number of endpoints contributing most to the variance between the two groups (Fig. 3B). Both alignment, characterized by a coherency value calculated by *TWOMBLI* that is analogous to the order parameter calculated by *AFT*, and number of matrix endpoints were significantly different between control and 21d βcat^istab^ mice (Fig. 3C). Specifically, 21d β-cat^istab^ mice had less alignment and a greater number of endpoints when compared to controls. Hyphal growth unit (HGU), which is a measure of the number of endpoints per unit fiber length, was also significantly lower in fibrotic mice (Fig. 3C). Similar to the data acquired using *AFT*, the fibrotic changes in collagen alignment, number of endpoints, and hyphal growth unit demonstrated by TWOMBLI were also rescued in dorsal dermis upon reversal from Wnt activation in 21d β-cat^istab^, 21r mice (Fig. 3E). Consistently, reversal mutants and litter matched controls did not separate considerably across any of the TWOMBLI matrix metrics (Fig. 3D).

One important consideration when capturing polarized light images of PSR-stained histological sections to be run through these image analysis algorithms is the angle of imaging [21]. Specifically, the aforementioned images used for AFT and TWOMBLI analyses were taken with the stage of the microscope at 0 degrees (Supplementary Fig. 1A). However, upon rotating the stage 45 degrees, different fibers were captured due to changes in the birefringence of the collagen fibers (Supplementary Fig. 1A, 1B). Consequently, different data was obtained when running these images through the image analysis algorithms, with different vector maps outputted by AFT (data not shown) and the separation across TWOMBLI metrics between 21d β-cat^istab^ mutants and littermatched controls lost in the PCA biplot (Supplementary Figure. 1C, 1D). Thus, it is important to select an angle that captures the fibers in the tissue of interest accurately when imaging with polarized light microscopy.

All together, our approach of using two different algorithms allowed us to analyze the topography of collagen at both the tissue and fiber levels. These results demonstrated that there are several topographical changes, including differences in collagen alignment, number of matrix endpoints, and hyphal growth unit during the onset of fibrosis. Using these algorithms, we were also able to determine that fibrotic changes in ECM topography can be rescued using our inducible and reversible Wnt activation model.

## Discussion

By using image analysis algorithms, we demonstrate that changes in collagen topography result from the onset of skin fibrosis. Specifically, there is a loss of collagen anisotropy, more matrix endpoints, and decreased hyphal growth unit in fibrotic skin. This is consistent with previous literature showing that during fibrosis, dermal thickening occurs due to fibroblast activation, resulting in increased ECM deposition [34]. With increased collagen fiber deposition, less organization and more endpoints are observed. Using our genetically inducible and reversible fibrosis model, we also showed that these changes in collagen topography can be reversed upon removal of a fibrotic stimulus. Here, we find the changes in collagen topography are accompanied by changes in collagen quantity and density, with an increase in dermal thickness and proportion of high density matrix during the onset of fibrosis that can also be rescued upon reversal [4]. Finally, we showed that these changes in collagen topography and quality can be quantified in an unbiased, efficient manner by adapting published image analysis algorithms.

In deciding which algorithm to use to examine the histomorphometry of fibrotic skin, we had three primary considerations. First, we determined that cell segmentation was not a priority. While segmentation allows for metrics to be calculated on an individual cell basis, PSR staining imaged using polarized light microscopy does not provide sufficient clarity for cell segmentation algorithms to accurately and reliably perform these calculations. Second, we had to consider differences between control and mutant samples, such as differences in intensity, which may alter metric calculations made by the algorithm. There are currently both intensity-independent and dependent software available. We tested the effect of intensity differences between each mutant group and its litter matched control on the utilized algorithms (data not shown). Finally, we had to consider the scale along which the algorithms calculate the histomorphometrics. Several metrics change drastically when evaluating on a few cells versus tissue scale, for instance [17]. Taking these considerations into account, we chose to use AFT and TWOMBLI, but other groups have conducted similar analyses using other softwares [19].

The fibrotic changes in collagen topography in our Wnt activation model have substantial implications for collagen quality and tissue integrity in skin and internal organs. The structural integrity and mechanical properties of a tissue is highly dependent on the topography of the ECM within it [15]. Maintaining collagen orientation within its homeostatic state is crucial for the structural integrity of the skin [15]. Specifically, collagen fibers in the skin typically follow a basketweave pattern to contribute to resistance of tensile stress in multiple directions [9]. Randomization of fiber orientation, as was observed in the dermis 21d β-cat^istab^ dorsal skin, may therefore contribute to the loss of tissue integrity that is observed during fibrosis [34]. Fiber orientation, in addition to the amount of high density matrix, contribute greatly to the mechanical properties of the skin, especially stiffness [15]. In connective tissue such as the skin, fibroblasts embedded within the ECM can sense the matrix stiffness caused by changes in fiber reorientation and density via heterodimeric transmembrane receptors called integrins [15]. This leads to the mechanoactivation of myofibroblasts via mechanotransduction pathways, such as YAP/TAZ and FAK, causing myofibroblasts to secrete more ECM and further propagate these topographical changes [11]. Ultimately, measuring topographical metrics at the tissue and fiber scale such as orientation provide a useful read-out for matrix and tissue integrity and mechanics.

Furthermore, while we applied image analysis algorithms to collagen, they can also be applied to other fibrous ECM components as well as cellular components. Fibrosis in skin and other organs such as lung are accompanied by lipodystrophy of adipocytes and lipofibroblasts [4, 35]. Activated fibroblasts are highly motile cells and contractile cells, as opposed to adipocytes, which are largely non-motile, resulting in each being characterized by a different organization of actin cytoskeleton [36]. Thus, examining the actin and cytoskeleton topography of these cells in fibrotic tissue using image analysis algorithms can indicate if there is a transition in early stages of fibrosis that can enable the change in morphology of fibroblasts and adipocytes to contractile cells which can also be contributors for elevated tissue stiffness.

Having widely-accessible and unbiased image analysis algorithms to analyze ECM topography has important implications for fibrosis research, clinical diagnosis, and treatment development [37]. Considering that there are currently no therapies to prevent or reverse existing fibrosis, image analysis algorithms can also be used to evaluate the success of potential therapies. This is especially true for therapeutics meant to target enzymes promoting matrix stiffening and mechanotransduction pathways, as ECM topography can be used as a read-out for changes in matrix stiffness and other aspects of matrix quality [11]. Together, this study shows that a very specific topography of ECM is present in the skin and this topography is altered during the onset of fibrosis. However, these alterations in ECM topography can be rescued upon removal of the fibrotic stimulus in our inducible and reversible Wnt activation model. Unbiased algorithms can be used to analyze the topography of ECM in fibrotic and other diseased tissues, providing new insights into tissue quality, cell behaviors, and reveal new therapeutic targets.

## List of Abbreviations

ECM: Extracellular Matrix
DWAT: Dermal White Adipose Tissue
AFM: Atomic Force Microscopy
PSR: Picrosirius Red
AFT: Alignment by Fourier Transform
TWOMBLI: The Workflow Of Matrix BioLogy Informatics
EnCre: *Engrailed 1* Cre-Recombinase
β-cat^istab^: β-catenin Inducibly Stabilized
FFT: Fast Fourier Transform
PCA: Principal Component Analysis
HGU: Hyphal Growth Unit
HDM: High Density Matrix

## Author Contributions

R.P.A. conceived, acquired funding for, and supervised the study; R.P.A. and R.H.W. designed experiments; R.H.W. and E.S. performed experiments; R.P.A. and A.R.J. provided resources and generated the tissues; R.H.W. adapted software; R.P.A. and R.H.W. analyzed data; R.H.W. and R.P.A. wrote the manuscript.

## Acknowledgements

We would like to thank all past and present members of the Atit Lab that contributed to this work. Special thanks to Sakin Kirti for technical support and advice. We thank the CWRU Imaging Core and bio for microscope use. This study is funded by National Institutes of Health-National Institute of Arthritis and Musculoskeletal and Skin Diseases R01 AR076938 (R.P.A.), National Institutes of Health T32 Musculoskeletal Predoctoral Training Grant T32 AR 7505-31 (A.R.J.), and the National Institutes of Health T32 Dermatology Predoctoral Training Grant T32 AR 7569-25 (A.R.J.). Schematics were made with biorender.com.

## Data Availability

Raw images can be made available upon request. Code available in GitHub (https://github.com/radhikaatit).

## Supporting Information

**Supplementary Fig 1.**
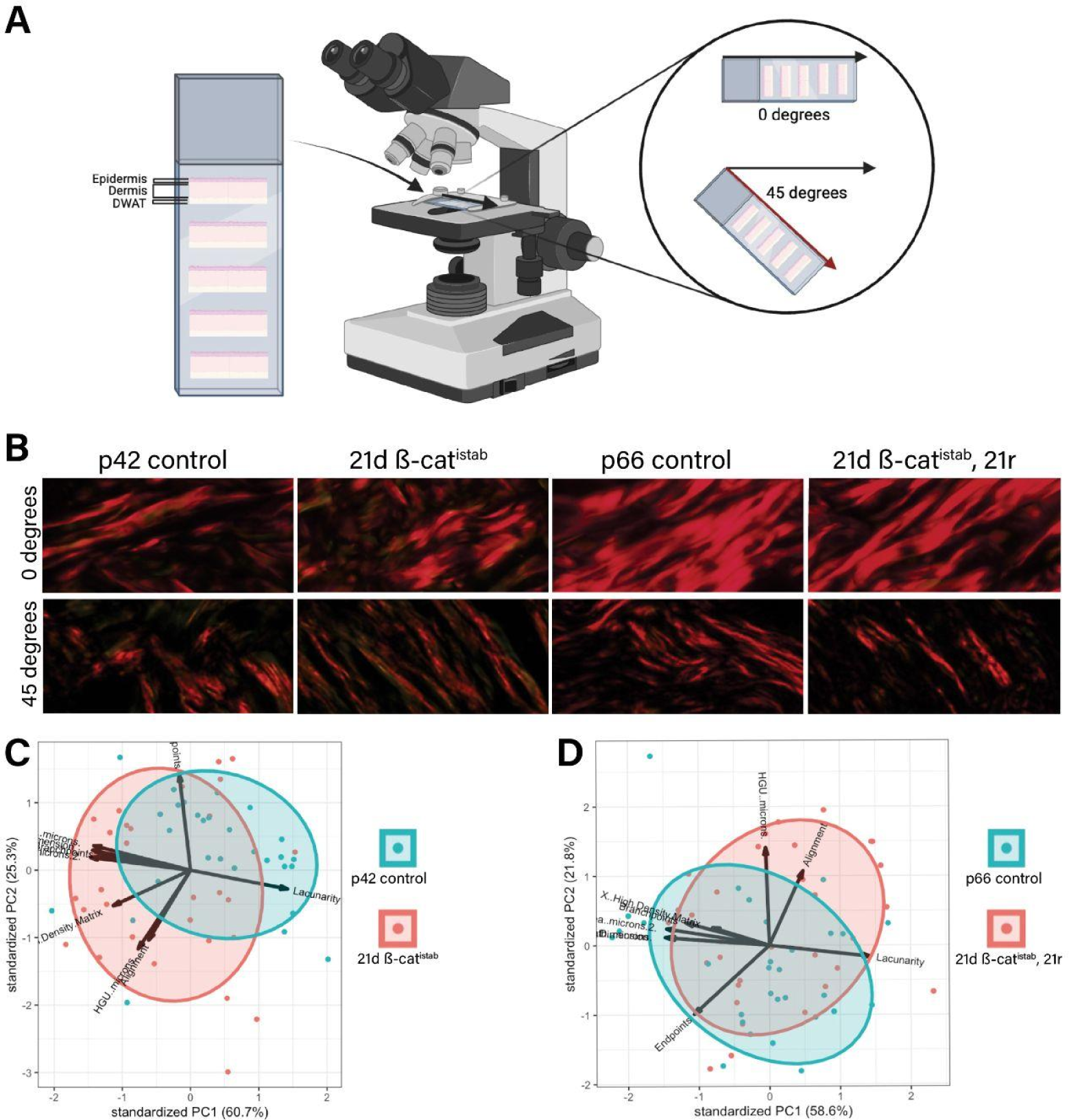
Angle of imaging influences PSR-stained collagen fibers detected by polarized light microscopy. (A) Schematic showing how skin sections were placed on charged slides and show slides were positioned to acquire images at 0 and 45 degree angles. (B) Cropped ROIs from 40x polarized light images of dorsal dermal collagen stained with picrosirius red at 0 and 45 degrees. (C,D) PCA plots showing the variance between each control/mutant pair and the TWOMBLI metrics that distinguish each control group and corresponding mutant group most when images were taken at a 45 degree angle. Each data point represents one ROI. 6 ROIs were used for each animal, and 5 animals were included for each condition.

